## Supplementary Material for "Crispant analysis in zebrafish as a tool for rapid functional screening of disease-causing genes for bone fragility"

### Supplementary figures:

**a**

| | *aldh7a1* | *daam2* | *esr1* | *sost* | *creb3l1* | *ifitm5* | *mbtps2* | *sec24d* | *serpinf1* | *sparc* |
|---|---|---|---|---|---|---|---|---|---|---|
| **InDelphi-mESC prediction** | 77% | 72% | 83% | 82% | 88% | 81% | 87% | 79% | 80% | 85% |

**b**

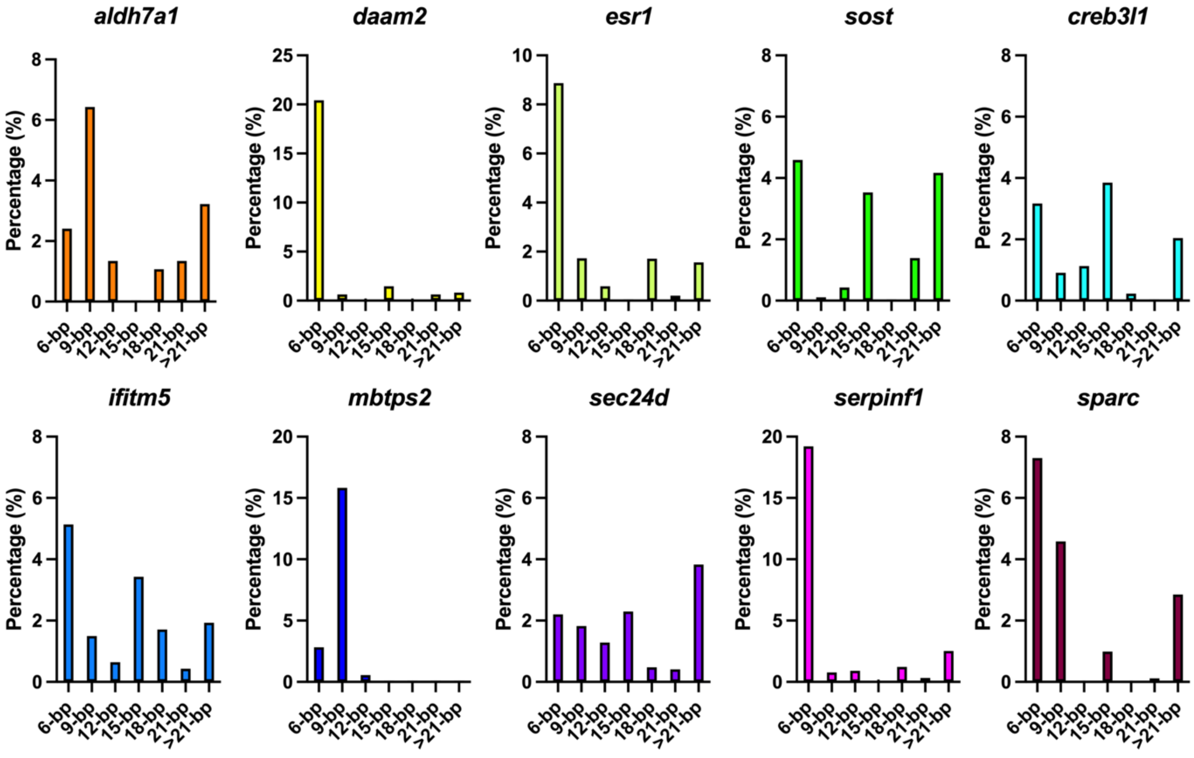


**Supplementary Figure 1: InDelphi prediction and In-frame analysis.** *The first four genes are associated with the pathogenesis of osteoporosis, while the last six are linked to osteogenesis imperfecta. (a) The predicted indel percentage using the InDelphi-mESC prediction tool. (b) Visualization of the in-frame analysis of the different crispants, showing the percentages of 9, 12, 15, 18, 21 and more than 21 base pair deletions in the crispants.*

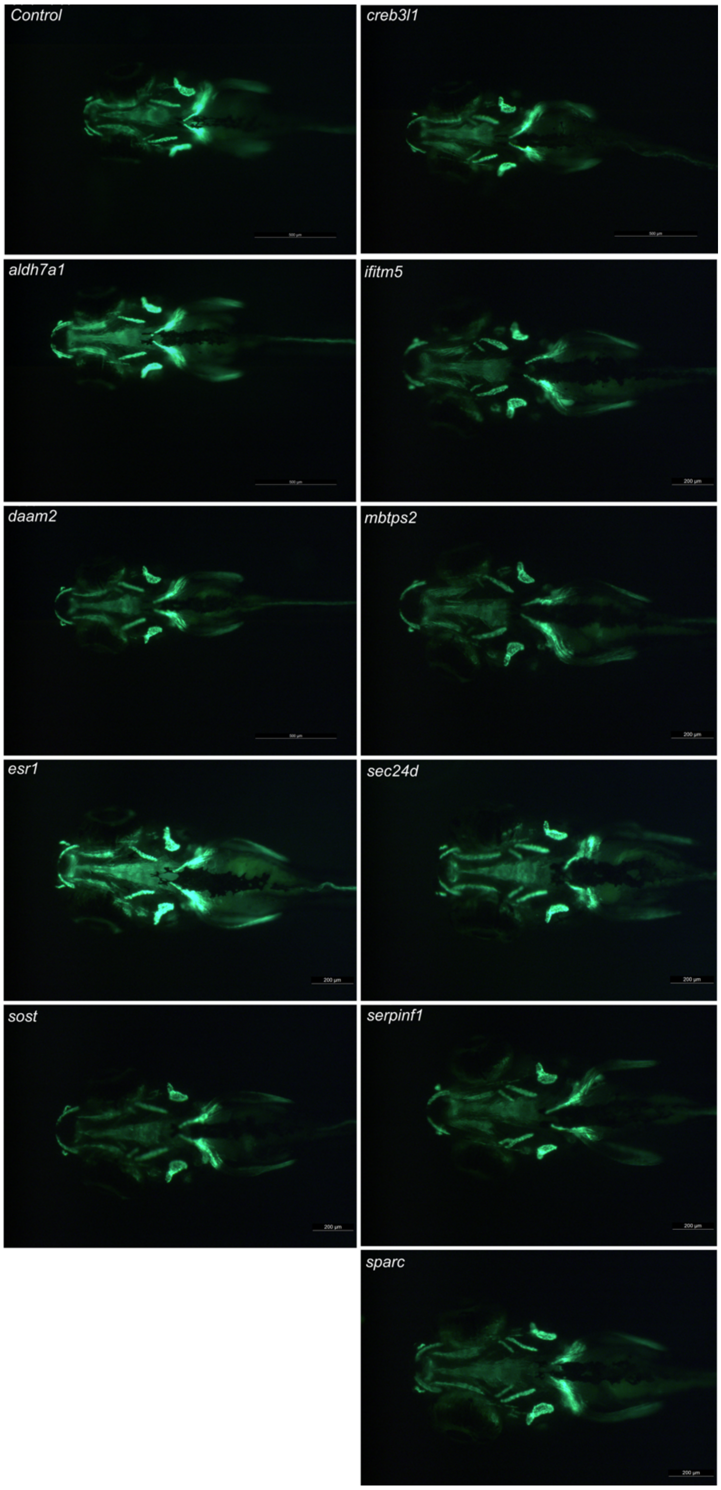


**Supplementary Figure 2: Osteoblast-positive head area at 7 dpf.** *Visualization of the osteoblast using the osx:Kaede transgenic line. The four genes on the left are associated with the pathogenesis of osteoporosis, while the six genes on the right are linked to osteogenesis imperfecta. The presented image shows a representative image of the specific crispants. Images are taken with the Leica microscope and the osx: Kaede positive larvae are visualized from a ventral perspective. Scale bars = 500 µm and 200 µm.*

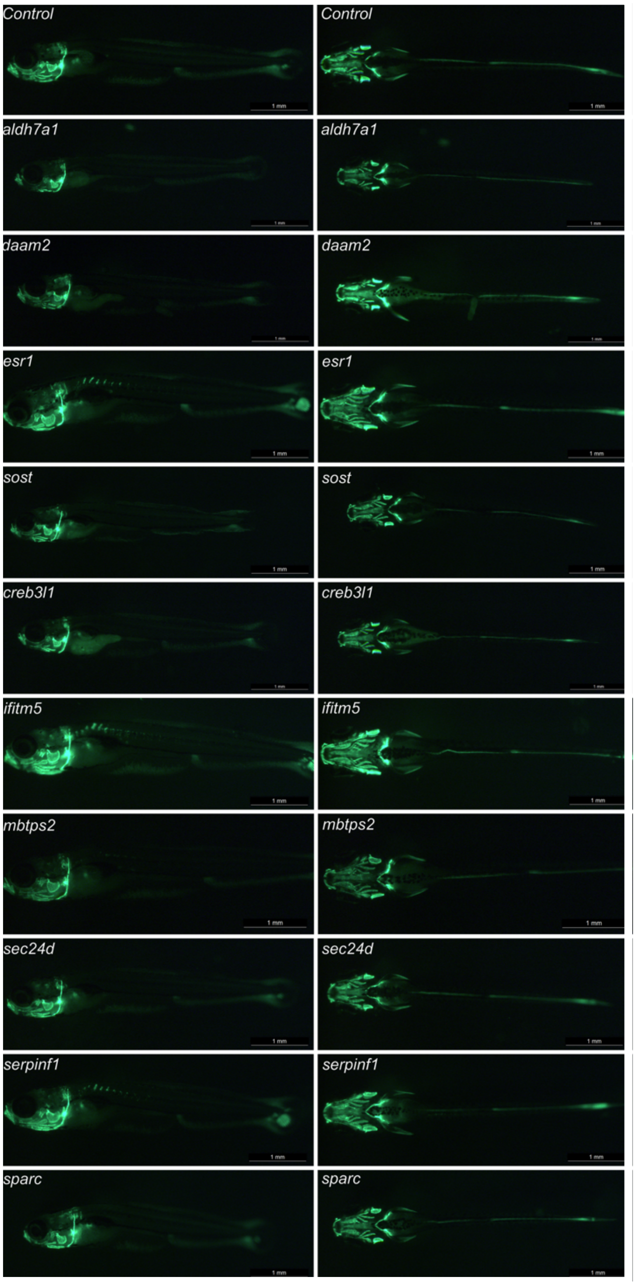


**Supplementary Figure 3: Osteoblast-positive head area at 14 dpf.** *Visualization of the osteoblast using the osx:Kaede transgenic line. The first four genes are associated with the pathogenesis of osteoporosis, while the last six are linked to osteogenesis imperfecta. The presented image shows a representative image of the specific crispants. Images are taken with the Leica microscope and the osx: Kaede positive larvae are visualized from a ventral and lateral perspective. Scale bars = 1 mm.*

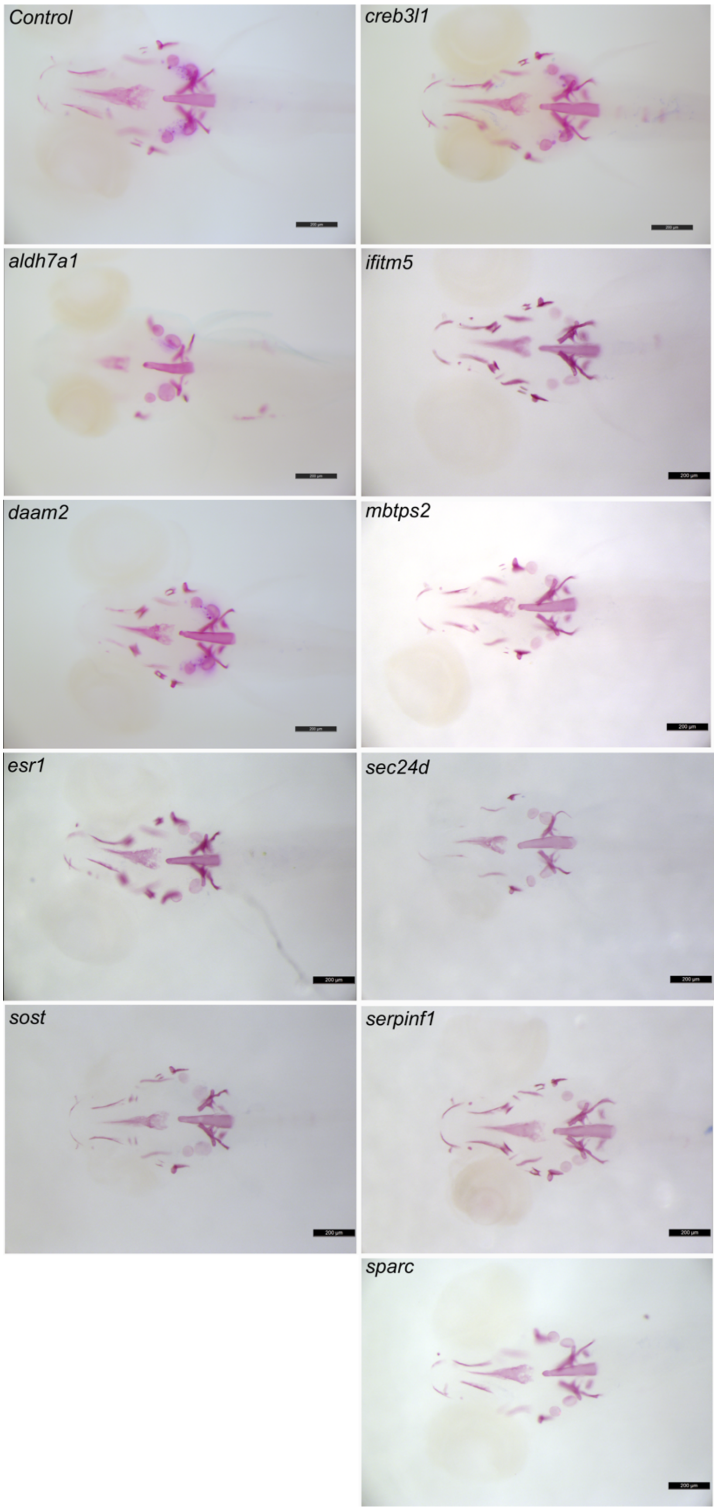


**Supplementary Figure 4: Mineralization in the head area at 7 dpf.** *Visualization of the mineralization after ARS staining. The four genes on the left are associated with the pathogenesis of osteoporosis, while the six genes on the right are linked to osteogenesis imperfecta. The presented image shows a representative image of the specific crispants. Images are taken with the Leica microscope and the stained larvae are visualized from a ventral perspective. Scale bars = 200 µm.*

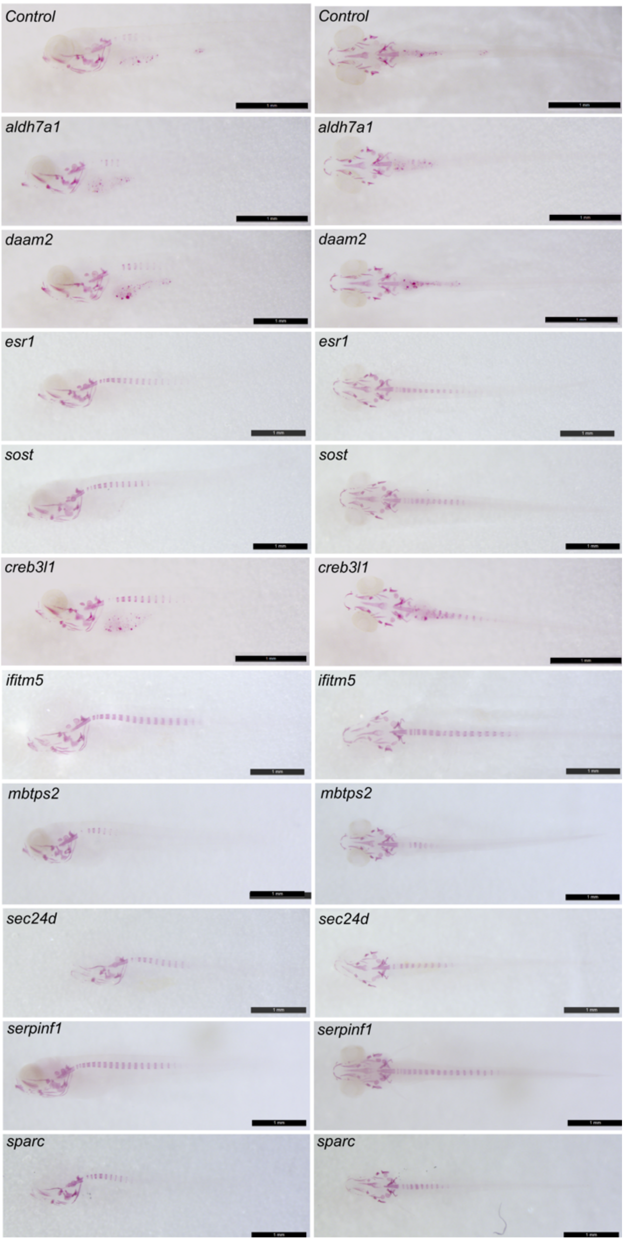


**Supplementary Figure 5: Mineralization in the head area at 14 dpf.** *Visualization of the mineralization after ARS staining. The first four genes are associated with the pathogenesis of osteoporosis, while the last six are linked to osteogenesis imperfecta. The presented image shows a representative image of the specific crispants. Images are taken with the Leica microscope and the stained larvae are visualized from a ventral and lateral perspective. Scale bars = 1 mm.*

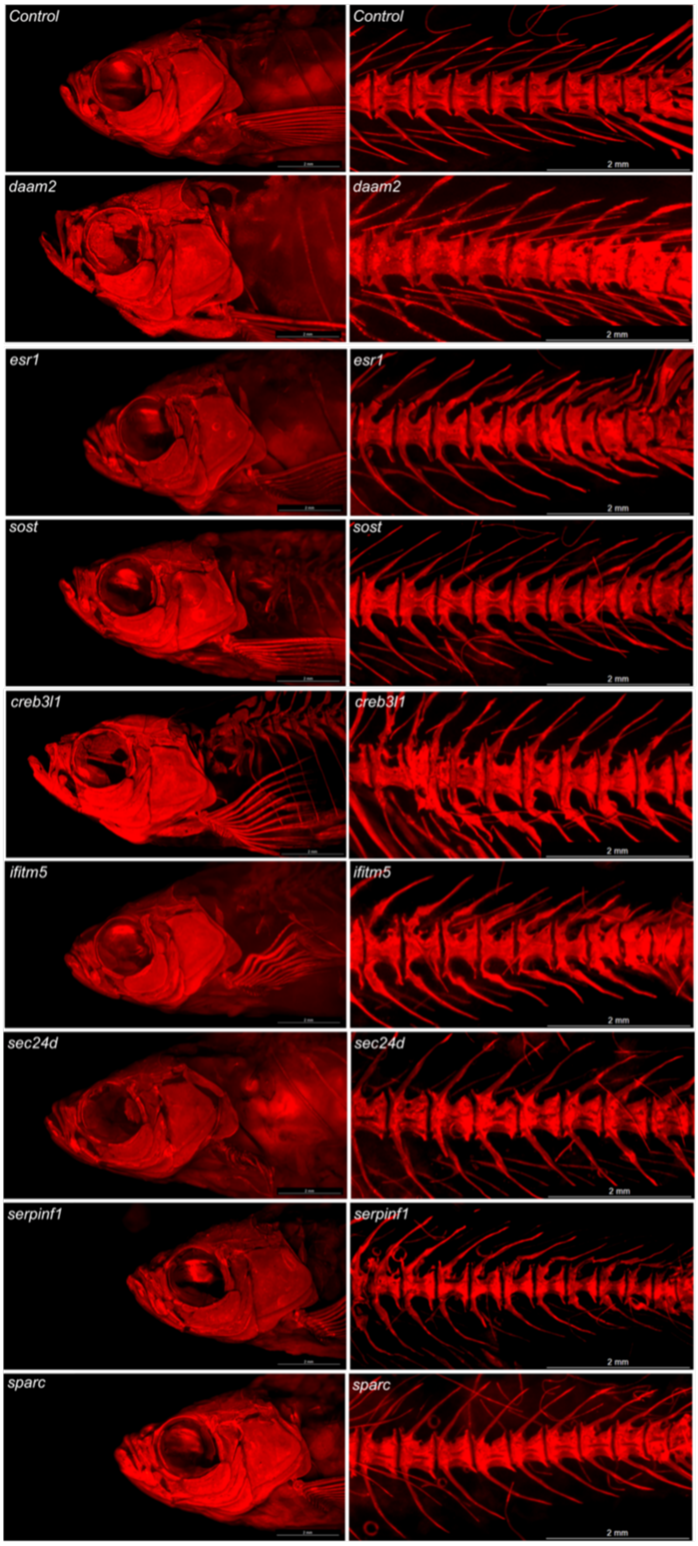


**Supplementary Figure 6: Mineralization in the skeleton at 90 dpf.** *Visualization of the mineralization after ARS staining. The first four genes are associated with the pathogenesis of osteoporosis, while the last six are linked to osteogenesis imperfecta. The presented image shows a representative image of the specific crispants. Images are taken with the Leica microscope and the stained adults are visualized from a lateral perspective. Scale bars = 1 mm.*

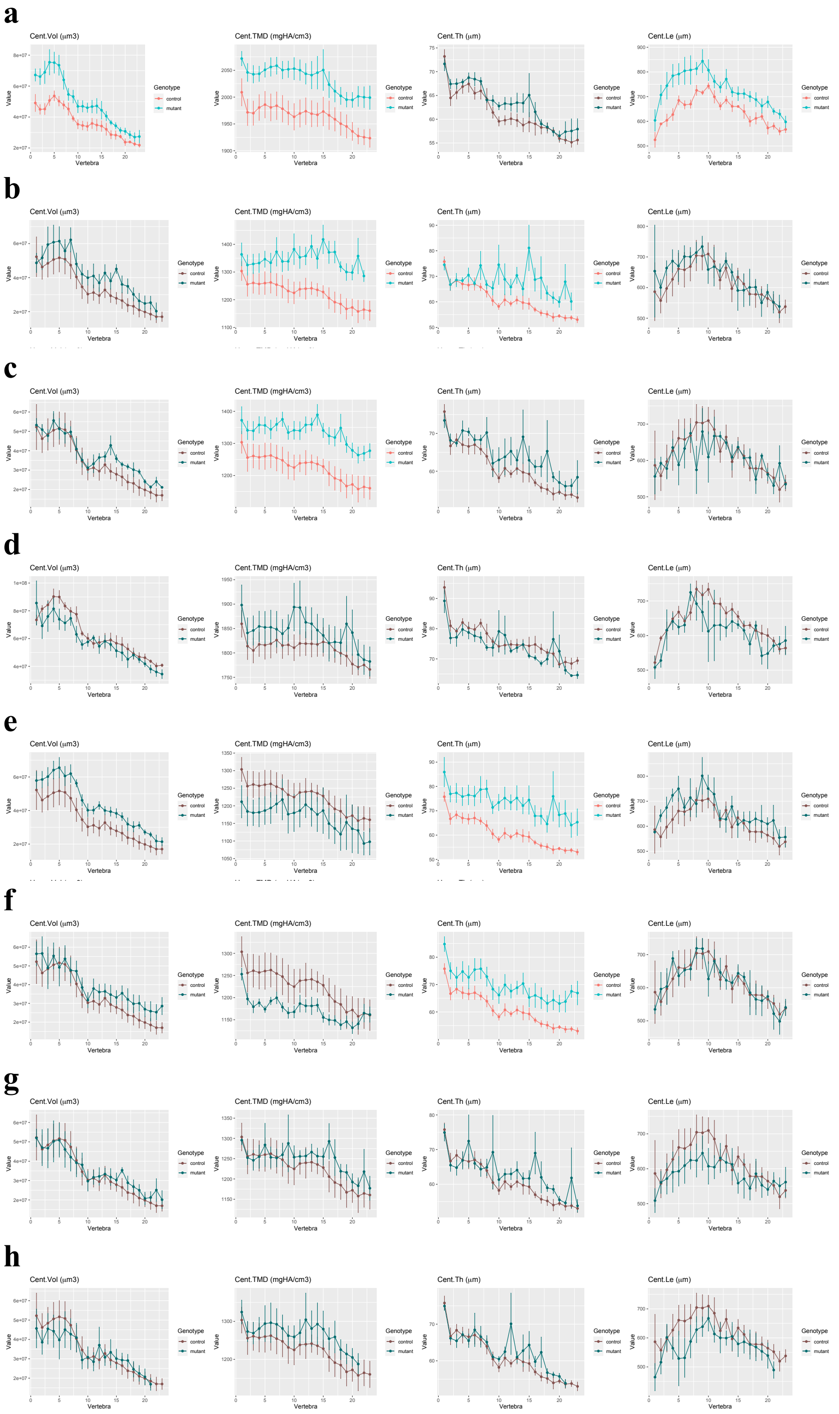


**Supplementary Figure 7: Quantitative µCT-scanning analysis of the vertebral column of the crispants using FishCuT software.** *Whole-body µCT-scanning was performed for the evaluation of skeletal structures in the vertebral column. Different bone related parameters are visualized: Tissue Mineral Density (TMD), Volume (Vol), Thickness (Th.), and Length (Le.) (a) Results of crispants for daam2. (b) Results for crispants for esr1. (c) Results for crispants for sost. (d) Results for crispants for creb3l1. (e) Results for crispants for ifitm5. (f) Results for crispants for sec24d. (g) Results for crispants for serping1. (h) Results for crispants for sparc. The first four genes are associated with the pathogenesis of osteoporosis, while the last six are linked to osteogenesis imperfecta. Statistically significant differences were represented with a lighter color scheme for easy visualization.*

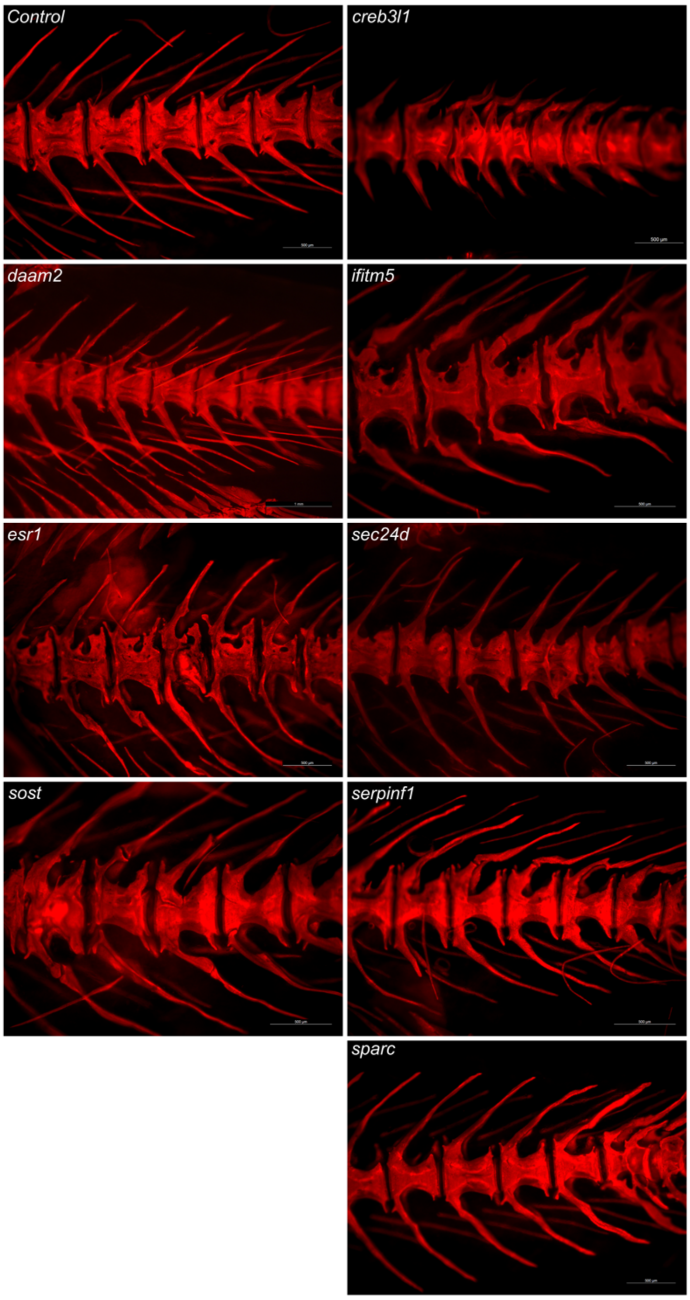


**Supplementary Figure 8: Mineralization in the skeleton at 90 dpf.** *Representative images of skeletal mineralization following Alizarin Red S (ARS) staining in a second clutch, demonstrating the consistency of the observed skeletal phenotype. The first four genes are associated with the pathogenesis of osteoporosis, while the last six are linked to osteogenesis imperfecta. Images show specific crispants from a lateral view, captured using a Leica microscope. Scale bars = 1 mm.*

**Supplementary Table 1:** *Overview of selected genes for crispant analysis, with reported mutations and/or polymorphisms associated with skeletal and non-skeletal phenotypes in human, mice and zebrafish. The conservation between human and zebrafish is reported in the last column. The first four genes are associated with the pathogenesis of osteoporosis, while the last six are linked to osteogenesis imperfecta.*

| | Human | | | Mouse | | | Zebrafish | | | |
|---|---|---|---|---|---|---|---|---|---|---|
| | Variant | Phenotypic features | Skeletal abnormalities | Mutation /model | Phenotypic features | Skeletal abnormalities | Mutation/model | Phenotypic features | Skeletal abnormalities | Conservation |
| *ALDH7A1* | c.1279G>C [1] | Pyridoxine - Dependent Epilepsy (PDE) phenotype | n/a | *Aldh7a1* KO mice [2] | Early death | n/a | *aldh7a1*-null zebrafish [3] | PDE phenotype with premature death at 14 dpf | n/a | 82% |
| | Polymorphisms [4] [5][6] | n/a | Associated with bone mineral density | / | / | / | *aldh7a1* Morpholino knockdown [7] | n/a | Defects in cartilage and pectoral fin development | |
| *DAAM2* | Missense mutation [8] | Nephrotic syndrome | n/a | Hypomorphic *Daam2* allele [9] | n/a | Reduced bone strength and increased cortical bone porosity | / | / | / | 72% |
| | Polymorphisms [9][10][11] | n/a | Associated with bone mineral density variation | / | / | / | / | / | / | |
| *ESR1* | Polymorphisms [12][13][14][15] [16][17][18] | Associated with different diseases, including breast cancer and | Associated with bone mineral density variation | *ERα* KO mice [19] | n/a | Decrease in cortical bone mineral density and increase in trabecular | Loss-of-function zebrafish [20] | Viable and overall normal Role of *esr1* in regulation of heart rate is discovered | n/a | 47% |

| | | | | | | | | | | |
|---|---|---|---|---|---|---|---|---|---|---|
| | | cardiovascular risk | | | | bone mineral density | | | | |
| *SOST* | Nonsense mutation [21] | Sclerosteosis | Progressive bone thickening and increased BMD | *Sost* KO mice [23] | n/a | Increase in bone volume, BMD and osteoblast surface area | n/a | n/a | n/a | 50% |
| | Polymorphisms [12][15][16][17][22] | n/a | Associated with bone mineral density variation | / | / | / | / | / | / | |
| *CREB3L1* | c.934_936delAAG [24]<br>c.1284C-A [25]<br>c.911C-T [26] | n/a | Osteogenesis Imperfecta type XVI | *Oasis* KO mice [27] | n/a | Decreased bone density, delayed osteoblast maturation resulting in severe osteopenia | *creb3l1*$^{\Delta bZIP/\Delta bZIP}$ zebrafish [28] | n/a | Upregulation of *creb3l1* expression | 59% |
| *IFITM5* | c. 14C>T [29]<br>c.119C>T [30]<br>c.119C>G [31] | n/a | Osteogenesis Imperfecta type V | c. 14C>T [32] | n/a | Severe skeletal defects: fractures and early lethality<br>Abnormalities in mineralization | n/a | n/a | n/a | 39% |
| | / | / | / | | n/a | | / | / | / | |

| | | | | | | | | | | |
|---|---|---|---|---|---|---|---|---|---|---|
| | | | | *Ifitm5* KO mice [33] | | No skeletal abnormalities | | | | |
| *MBTPS2* | Missense mutation [34][35] | X-linked Olmsted syndrome and IFAP Syndrome 1, with or without BRESHECK Syndrome | X-linked Osteogenesis Imperfecta | *Mbtps2* Knock-in (N455S) [36] | n/a | Embryonic lethality in hemizygous male mice, early osteoarthritis in heterozygous female mice | n/a | n/a | n/a | 60% |
| | / | / | / | *Mbtps2* KO mice [36] | n/a | Embryonic lethality in hemizygous male mice, early osteoarthritis in heterozygous female mice | / | / | / | |
| *SEC24D* | Nonsense variants [37][38] Missense variants [37][38][39] Frameshift variants [40][41][39] | n/a | Autosomal recessive Osteogenesis Imperfecta with a Cole-Carpenter syndrome-like phenotype | *Sec24d* KO mice [42] | Embryonic lethality | Embryonic lethality | *sec24d* KO zebrafish (*Bulldog*) [43] | | Abnormalities in pectoral fin and head skeleton | 65% |

| | | | | | | | | | | |
|---|---|---|---|---|---|---|---|---|---|---|
| *SERPINF1* | c.696C>G [44]<br>c.324_325dupCT [44]<br>c.1132C>T [44]<br>c.1118_1119del [45]<br>c.1-4796dupT [45]<br>c.653delT [45] | n/a | Osteogenesis Imperfecta type VI | PEDF KO mice [46] | Retinal malformations | Mild reduction in trabecular bone volume and accumulation of unmineralized bone matrix: increased bone fragility | n/a | n/a | n/a | 39% |
| *SPARC* | c.497G>A [47]<br>c.787G>A [47] | n/a | Osteogenesis Imperfecta type XVII | *Sparc* null mice [48] | Abnormal eye phenotype | Decrease in BMD, bone mineral content and increase in bone fragility | n/a | n/a | n/a | 77% |

**Supplementary Table 2: Assessment of off-target effects in crispants.** *Genes that are possibly targeted by the selected crRNA for each of the crispants are called 'off-target genes' in this table (mm=number of mismatches). The top 3 ranked off-target effects, selected based on a high CFD (cutting frequency determination) score, are listed, together with their chromosomal position, forward and reverse primer for amplification, the CFD (cutting frequency determination) score and the off-target percentage, based on NGS analysis of a pool of DNA of 1-day old crispants (n=10).*

| Information off-target gene | Chromosomal location (GRCz11) | Forward primer | Reverse primer | CFD score | Off-target percentage |
|---|---|---|---|---|---|
| | | *aldh7a1* | | | |
| mm4_intergenic_pcdh1b\|rnasel3 | Chr.14: 38013913 | TGGACCAGCCTGCCTATTAT | GAGCCCATTCCTCTCATAGC | 0.35 | 0% |
| mm3_intron_cdh2 | Chr.20: 17740399 | GAGCGATGCTGCTGCTCTAT | CGAACAGAAACCCGACAGGA | 0.23 | 0% |
| mm4_intergenic_si:ch73-321d9.2\|ndufc2 | Chr.15: 1139538 | GGGGAAAGTGCCTAGGCTTT | GGGGTGGACTAAGGCAAGTG | 0.19 | 0% |
| | | *daam2* | | | |
| mm4_intron_zdhhc8b | Chr.5: 18002939 | GCAAAACGATCTCTGTGCAC | TGCCTACGATGTTGTTGGAC | 0.54 | 0% |
| mm4_intergenic_CR388008.1\|ppp2r2ab | Chr.10: 18577094 | CCTTGTTCAAAGCTTCCTGC | TCACATGACATACCAGCACA | 0.35 | 1,20% |
| mm4_intron_klc1a | Chr.13: 15820767 | TGTTACCCTTGCTTTGCCAT | CAAGCATACAAGCATAGCGC | 0.33 | 2,82% |
| | | *esr1* | | | |
| mm4_intron_cyp11a1 | Chr.25: 22329136 | GCACGGACAGCTACTTTTGC | GGTGGACATTCCTGCAGTCA | 0.26 | 0,11% |
| mm4_intron_ctnna1 | Chr.24: 34548469 | CGAGATGGAACCAAATGAGTGC | CCTCTCTGGCCATGCACTAC | 0.26 | 0,60% |
| mm4_intron_prmt3 | Chr.7: 17170999 | ACGGTCAAGGCTGCCTTTTA | TCTCAAGCATCAGACTTGGAGT | 0.23 | 0,26% |
| | | *sost* | | | |
| mm4_intergenic_zbtb12.1\|zbtb12.2 | Chr.19: 26947788 | TTTCGGCCATATCTGCCCAA | CAATCGTAGCTCGCTGTTGC | 0.50 | 0% |
| mm4_intergenic_RF00001\|rapgef2 | Chr.14: 47605837 | GCAAGCACATCTTCTCAGCC | TGACAGGTTTCTCCCAAACGA | 0.33 | 1,56% |
| mm4_intergenic_aim1a\|FP015823.1/aim1a | Chr.17: 25488841 | TCCTCTTTCATTCAGATCCATGA | ATGGTGTCTTGGCGGCATAA | 0.23 | 0,20% |
| | | *creb3l1* | | | |
| creb3l1_mm4_intergenic_dlg1l\|ptmab | Chr.2: 5109921 | TTGAGTTTCCATGGGCTGCA | GGGAGAGAGAGAGACGAGCA | 0.42 | 1,37% |
| creb3l1_mm4_intron_slc38a3a | Chr.11: 34832596 | TCGAAGGGCCCCTAGTACAA | TCATGCCAGGCCAGTAGATG | 0.31 | 0% |
| creb3l1_mm3_intergenic_ciartb\|fbxl22 | Chr.25: 26709599 | GGGTAGATCAGGGTCACACG | ACCCAGAGAATTAGGACGGC | 0.28 | 0% |
| | | *ifitm5* | | | |
| mm4_intergenic_nicn1\|FO904977.1 | Chr.22: 33524479 | TGGCCTTGATGAATGCGGAT | ATGCCATAGACACCACAGCT | 0.57 | 0% |
| mm3_exon_mylk4b | Chr.20: 26865816 | GTGCAGAAGAGCGACGGATA | CTGAGTACGCGACCTGCTAG | 0.41 | 0% |
| mm3_intron_col18a1a | Chr.9: 41944389 | CCAGGCCATTTCCAGAGCAG | GCAGTCCACACCTTACAGCC | 0.14 | 0% |
| | | *mbtps2* | | | |
| mm4_exon_BX088696.1 | Chr.8: 45725913 | TGGTGTTGTAAGTGGTGGTG | CATGCTGCTAACCGTTGTTG | 0.68 | 0% |
| mm4_intron_zgc:194007 | Chr.8: 42713866 | AGTTTTCTGTTGTGGTAGAGCA | CCATGGACGCAGAGCAATAT | 0.34 | 0% |
| mm4_exon_abcb10 | Chr.13: 24266797 | TGCAGTGATTCTGGTGTCAA | GATGTCTTACCTGTGTGGCC | 0.16 | 0% |
| | | *sec24d* | | | |
| mm3_intron_ntng2b_chr21_4021964_R | Chr.21: 4021978 | AGCAGCTAGGGGATCAGACT | TGTCCGTGTTTTCTCTCCCC | 0.16 | 0% |
| mm3_exon_obscnb | Chr.24: 35975523 | TTGACAAGCCAGGAGGTCAC | TACTCGCCAGCATCTTCCAC | 0.14 | 0% |
| mm4_intergenic_smim19\|egr1 | Chr.14: 21327480 | TGTTACACGGCTGTCTGAGA | TGACCGAATGAAGCTGATGAA( | 0.03 | 0% |
| | | *serpinf1* | | | |
| mm4_intergenic_BX569787.1/nr2f2\|nr2f2 | Chr.18: 23945771 | AGGGCCAAGGAGAGGAGATT | TTTGCCTCCATCCCGTTTCA | 0.47 | 0% |
| mm4_intergenic_BX664603.1\|mad2l1 | Chr.7: 57634995 | TGACGTTGAGAACAGATCAAACA | TCTTGTTGCTCTGCCTCTGG | 0.42 | 0% |
| mm4_exon_fus | Chr.3: 32701042 | ATGGACAGTCCCAGTCAGGT | GAATTCTCACCCAGGGGCAG | 0.18 | 2,13% |
| | | *sparc* | | | |
| mm4_intergenic_RF00001\|BX322647.1 | Chr.1: 33083559 | CACAGCGGCAAGTCAATGAG | GCGCAAAGTGCACACTACAA | 0.39 | 0% |
| mm4_exon_si:dkey-256e7.5_chr1 | Chr1.: 17668323 | ATCTGCTTAGCTGTGACGGG | TGGAAGTTGAGTTGAAGGAGC/ | 0.28 | 0% |
| mm4_intergenic_CU929161.1\|RF00026 | Chr.1: 7111923 | GGTATCAGCCGAAGTCGAGG | GTGTCAGGACAAGAGGAGCA | 0.14 | 0,63% |

**Supplementary Table 3: Crispant genotyping.** *Crispant genes with crRNA sequence, forward and reverse primers and assay specifications for Next-generation sequencing (NGS) are listed in this table. The first four genes are associated with the pathogenesis of osteoporosis, while the last six are linked to osteogenesis imperfecta. Primers are designed using Primer3 (https://primer3.ut.ee).*

| Gene | crRNA | Forward primer | Reverse primer | Assay |
|---|---|---|---|---|
| *aldh7a1* | AATTGTTCGACAGATTGGAG | CCTATTTGCTTAGTTAAATTCAGGTGG | CATCCACATACTCCTGCACC | FORD58 |
| *daam2* | GATCTATTGCAGTAAGAAGA | AGTGTGGCTCAGTTGATGTGT | ACATGTCTTTGGATTGCCCCT | FORD60 |
| *esr1* | CAGCTCCTCACACAGGCCCA | TGTTTCCATGGTGATGTCTGG | TGTTAGTAAATACATCAACTCAAAGACC | FORD60 |
| *sost* | GTTTGATCACAATATGCCAG | GTGCCGTCCATCAACAGC | GGTGAACTTGAAATGAACCGC | FORD60 |
| *creb3l1* | ACACAGTTACTCTCTCAGCG | TGTATCCGTTTGCTACCATACC | TGTTTGTGTGGATGGATGGC | FORD60 |
| *ifitm5* | AGCGCAGGAATGCTCAGACA | TCCATCAAGGCTCGAGATCAG | TGAGAAATTCACACAGCCAAGG | FORD60 |
| *mbtps2* | TTCCATATCAAGTGGCACAC | TCTCCTTCTGTCTGTTTCACC | ACTCGGATGGAAAGCAAAGC | FORD60 |
| *sec24d* | GCCTATGGATCTCCAACACA | AAATTGAAACTACAGTGCATAATGG | GGGTCCATTGTTCATCTGAGG | FORD60 |
| *serpinf1* | GTGTTACGAGGAAGAGGGGA | TCTCTCTGTCATTCCGCTGG | GTAAATCTGCTTCTCGGCCC | FORD60 |
| *sparc* | CTGCCAGAGTCTTGCCAGCG | CTACAGCCCTTTCATTCCAGG | TGACCCAAACACATAACTATAAGC | FORD60 |

**Supplementary Table 4: qPCR primers.** *Skeletal marker genes and reference genes with forward and reverse primers are listed in this table. Primers are designed using NCBI PrimerBlast ([https://www.ncbi.nlm.nih.gov/tools/primer-blast](https://www.ncbi.nlm.nih.gov/tools/primer-blast)).*

| Gene | FWD primer | RVS primer |
|---|---|---|
| | **Skeletal marker genes** | |
| *runx2a* | GGAAGAGGAAAGAGCTTCAC | CGTCCACTGTGACCTTTATG |
| *sp7* | CTCTCCTCTCCCGCTTT | GTGTTTCCTCCTCCAGAATC |
| *sox9a* | TGGGAAAACTTTGGAGATTACTGA | AGTCGGGGTGATCTTTCTTGT |
| *sox9b* | GAAGATGGAGAGCAGACGCA | CCTGAGACTGACCGGAGTG |
| *bglap* | TCTTCCTGACTCCTCAGATAC | AGCCCTCTTCTGTCTCAT |
| *col1a1a* | TCTGGTGGCTTTGATGAG | GGGACCAGTAAATCCTGGG |
| *col1a1b* | CTTGCAGTGAGAGGACAA | GCTCGGGTTTCCATACAT |
| *col1a2* | TAACCCTGGTGCTAATGGTA | ACACCAGAATCTCCCTTCA |
| *col2a1a* | GAGAACCAGGCGATATTACA | CACCCTTAGCTCGTCTTTC |
| *col2a1b* | GTTGGTACAGCAGGATCTC | TCCAGGAACACCAGACT |
| | **Reference genes** | |
| *elfa* | GGAGACTGGTGTCCTCAA | GGTGCATCTCAACAGACTT |
| *bactin2* | ACGATGGATGGGAAGACA | AAATTGCCGCACTGGTT |
